# The sphingolipid inhibitor myriocin increases susceptibility of *Candida auris* to amphotericin B

**DOI:** 10.1101/2022.04.14.488279

**Authors:** Hanna Stieber, Lara Junghanns, Hannah Wilhelm, Maria Batliner, Alexander Maximilian Aldejohann, Oliver Kurzai, Ronny Martin

**Affiliations:** Institute for Hygiene and Microbiology, University of Würzburg, Würzburg, Germany; National Reference Center for Invasive Fungal Infections, Leibniz Institute for Natural Product Research and Infection Biology- Hans Knoell Institute, Jena, Germany; Research Group Fungal Septomics, Leibniz Institute for Natural Product Research and Infection Biology- Hans Knoell Institute, Jena, Germany

**Author notes:** Corresponding author: Ronny Martin, University of Würzburg, Institute for Hygiene and Microbiology, Josef-Schneider-Str. 2 / Building E1.

**Keywords:** *Candida auris*, myriocin, amphotericin B, sphingolipids, antifungal drug resistance

## Abstract

The emerging human fungal pathogen *Candida auris* can cause of hospital outbreaks by transmission between patients and / or medical care staff. It is known for a fast adaption to environmental stress and antifungal treatment. Besides resistance to fluconazole in most isolates, *C. auris* frequently shows limited susceptibility to echinocandins and even to amphotericin B. Therefore, multidrug resistance (MDR) is frequently found in clinical isolates, leading to a need for new therapeutic targets and compounds. One of those targets could be sphingolipids which are an integral part of the cell membrane and are important for membrane fluidity and the formation of lipid rafts. Here, we show that myriocin-mediated inhibition of the *de novo* sphingolipid biosynthesis caused severe growth defects in *C. auris*. Sublethal myriocin concentrations increase fungal susceptibility to amphotericin B and anidulafungin. While the effect is limited for anidulafungin, even isolates that show phenotypic resistance to amphotericin B became susceptible in presence of myriocin. A large-scale screening of 59 clinical isolates validated these initial results and showed that inhibition of the *de novo* sphingolipid biosynthesis increases *C. auris* susceptibility to amphotericin B. These observations open new options for future therapeutic strategies and illustrate that sphingolipids might play a pivotal role for the ability of *C. auris* to develop resistance against amphotericin B.

## Introduction

The human pathogenic yeast *Candida auris* was first discovered as a causative agent of external otitis in a Japanese patient (1). It is a close relative of *C. haemulonii, C. pseudohaemulonii* and *C. duohaemulonii* and together they form a cluster within the CTG clade which also contains medically important species like *C. albicans, C. tropicalis, C. parapsilosis, Meyerozyma guilliermondii* and *Claviceps lusitaniae* (2). *C. auris* is haploid and the species can be divided into four to five geographically distinct clades (3-5). Since its identification, *C. auris* emerged from a curiosity to a globally important human fungal pathogen which is characterized by high resistance to environmental stresses and antifungal drugs (6-10).

Another major clinical problem is the easy transmission of *C. auris* between patients and care personal, frequently causing outbreaks in medical care facilities (3, 11-14). Nearly all clinical isolates showed an intrinsic fluconazole resistance and high minimal inhibitory concentrations for echinocandins and even amphotericin B (3, 13, 14). Especially clade I isolates often developed MDR while those from clade II were the most susceptible ones (14). The frequent findings of isolates with resistance against two or even all three antifungal classes limit therapeutic options. In addition, *C. auris*-specific breakpoints for antifungals are not defined yet. Previous approaches to meet these challenges by identifying new therapeutics included large-scale compound testing and predatory yeasts (15, 16).

As cell membrane components, sphingolipids are involved in a variety of biological processes of the fungal cell and might be an interesting target for new treatment approaches (17). Changes of the sphingolipid composition are reported to influence susceptibility of *Candida* species to azoles, echinocandins and amphotericin B (18-21). Recent works have already shown that the composition of sphingolipids in *C. auris* is different from other *Candida* species (22-24). For example, glycosylceramides are the dominant sphingolipid species in *C. auris*, but absent in *C. glabrata* (15,25). Interestingly, *C. auris* was found to be highly susceptible to the sphingolipid biosynthesis inhibitor myriocin (15, 24). Based on these observations, we tested the susceptibility of different clinical *C. auris* isolates to myriocin and consequently examined the effects of a combination of the inhibitor together with antifungal drugs. We could show that presence of myriocin increases the susceptibility of *C. auris* to amphotericin B and partially to anidulafungin but not to fluconazole.

## Results

### *C. auris* more susceptible to myriocin than other *Candida* species

Initially, several medically important *Candida* species were grown on YPD medium with 1 and 5 µM myriocin or methanol which was used as solvent control. After three days of growth at 37°C, we observed that only *C. auris* displayed dramatic growth defects already at 1 µM myriocin, while all other Candida species exposed to this concentration were not affected (Figure 1 A). Aside *C. auris, C. glabrata* and *C. parapsilosis* did not show growth at 5 µM. A reduction was observed for *C. albicans* and *C. dubliniensis*. Interestingly, *C. tropicalis, Claviceps lusitaniae* and *Meyerozyma guilliermondii* were highly resistant against myriocin as their growth was not impaired at 5 µM (Figure 1 A). Due to these observations, we continued to analyze the susceptibility of *C. auris* B8441, *C. albicans* SC5314 and *C. glabrata* CBS138 on the rich RPMI1640 medium which had a pH of 7.3 and the minimal medium SDG which only contained glucose and yeast nitrogen base. On both media severe growth defects for *C. auris* in the presence of 1 µM and 5 µM myriocin were detected, although they were more pronounced on SDG medium (Figure 1 B). In presence of myriocin, *C. glabrata* grew better on RPMI1640 and SDG than on YPD (Figure 1).

**Figure 1.**
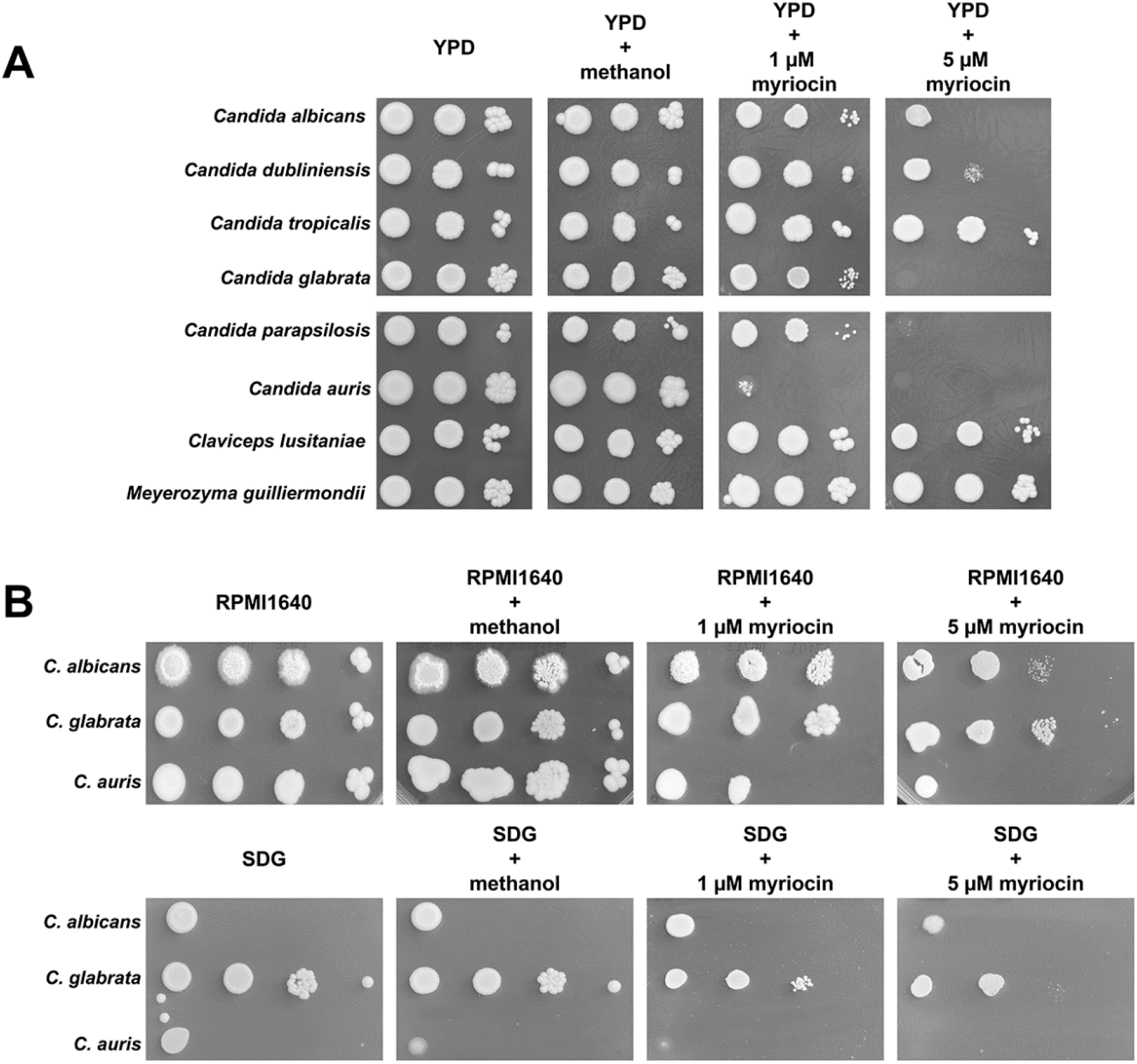
*C. auris* is highly susceptible to the sphingolipid inhibitor myriocin. (A) Several medically important *Candida* species were grown on YPD medium with or without different myriocin concentrations for 3 days at 37°C. Methanol was used as solvent control for myriocin. *Candida* cell concentrations are from left to right: 1x 10^6^ cells/ml, 1x 10^4^ cells/ml and 1x 10^2^ cells/ml. (B) *C. albicans* SC5314, *C. glabrata* CBS138 and *C. auris* B8441 were grown on RPMI or SDG medium with or without different myriocin concentrations for 3 days at 37°C. Fungal cell concentrations were 1x 10^8^ cells/ml, 1x 10^6^ cells/ml, 1x 10^4^ cells/ml and 1x 10^2^ cells/ml. from left to right.

### Myriocin strongly increases amphotericin B susceptibility

As *C. auris* is known for its high grade of resistance and / or tolerance to antifungal drugs, we further examined if the addition of myriocin would have an impact on the fungal susceptibility against fluconazole, anidulafungin and amphotericin B. They represent the antifungal drug classes of azoles, echinocandins and polyenes. Based on the initial observations, we tested the antifungal drug susceptibility with or without myriocin concentrations staying below 1 µM avoiding growth deficiencies. The Tests were performed with *C. auris* reference strain B8441 which is regarded as resistant to fluconazole and susceptible to anidulafungin and amphotericin. E-tests for fluconazole, anidulafungin and amphotericin were applied to RPMI1640 plates containing 2.5 to 250 nM myriocin and cells of *C. auris* B8441. As can be observed for medium only and the solvent control methanol, this strain is resistant to fluconazole (Figure 2). However, we observed an increased susceptibility and a MIC decrease from more than 265 µg/ml to 0.75 µg/ml in presence of 250 nM myriocin (Figure 2). However, only high doses of the sphingolipid inhibitor increased fungal susceptibility to fluconazole whereas no effects were detected for lower myriocin concentrations (Figure 2). Although B8441 has no FKS hotspot mutation, the strain has a MIC of 2 µg/ml for anidulafungin. In presence of myriocin, this MIC decreased to 0.25 µg/ml, even with low concentrations of the inhibitor (Figure 2). The increase of the fungal susceptibility became more pronounced with increasing myriocin concentrations (Figure 2). *C. auris* B8441 had an MIC for amphotericin B of 0.25 µg/ml on RPMI medium alone or with methanol (Figure 2). Strikingly, presence of 250 nM myriocin decreased the MIC to 0.023 µg/ml and an increase of the inhibition zone diameter (in comparison to no myriocin) was observed (Figure 2). These effects were still visible but less pronounced for 25 nM and 2.5 nM myriocin (Figure 2). In summary, inhibition of the sphingolipid biosynthesis by myriocin increases the fungal susceptibility to echinocandins and polyenes.

**Figure 2.**
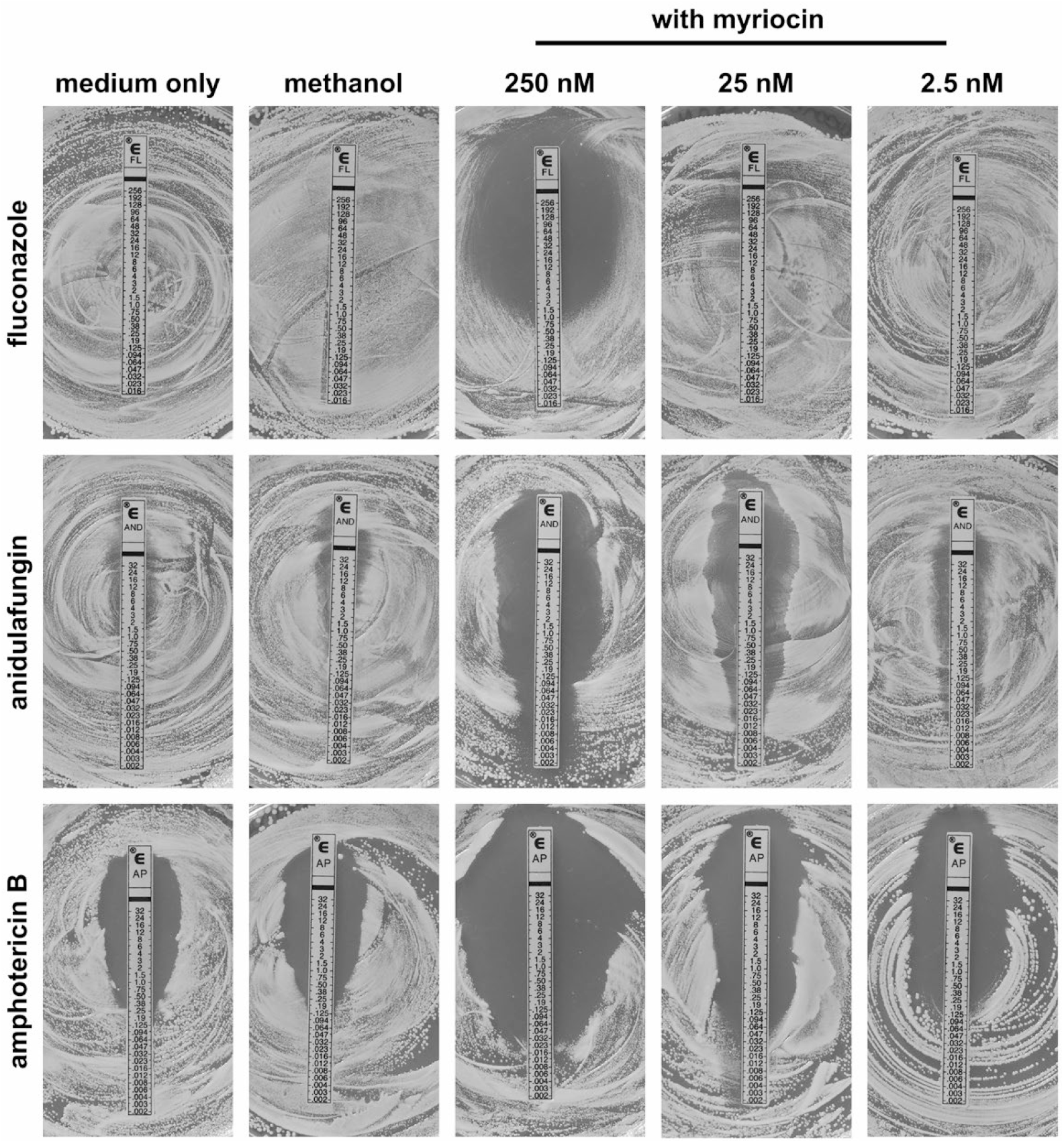
Addition of myriocin increases antifungal drug susceptibility of *C. auris* reference strain B8441. Cells of the *C. auris* strain B8441 were plated onto RPMI1640 agar with or without the indicated concentrations of myriocin or methanol as solvent control. The indicated E-tests were applied, and plates were incubated at 35°C for 48 h before pictures were taken.

### Combination of myriocin and antifungals had different effects on resistant strains

Encouraged by these observations, we further examined myriocin effects on two isolates which were previously found to be resistant to resistant against fluconazole and one of the other two antifungals. *C. auris* NRZ-2017-505 is resistant against echinocandins and has a verified *FKS1* S639Y mutation (26). Presence of 2.5 to 250 nM myriocin did not alter the susceptibility of the strain against fluconazole and anidulafungin as the MICs remained beyond 256 µg/ml for fluconazole and 32 µg/ml for anidulafungin (Figure 3). In contrast, myriocin increased susceptibility to amphotericin B, leading to a decrease of MIC-values from 1 µg/ml to 0.047 µg/ml in the presence of 250 nM myriocin and to 0.38 µg/ml in the presence of 25 nM myriocin (Figure 3). A concentration of 2.5 nM had no effects as the MIC for amphotericin remained at 1 µg/ml as for RPMI medium alone (Figure 3).

**Figure 3.**
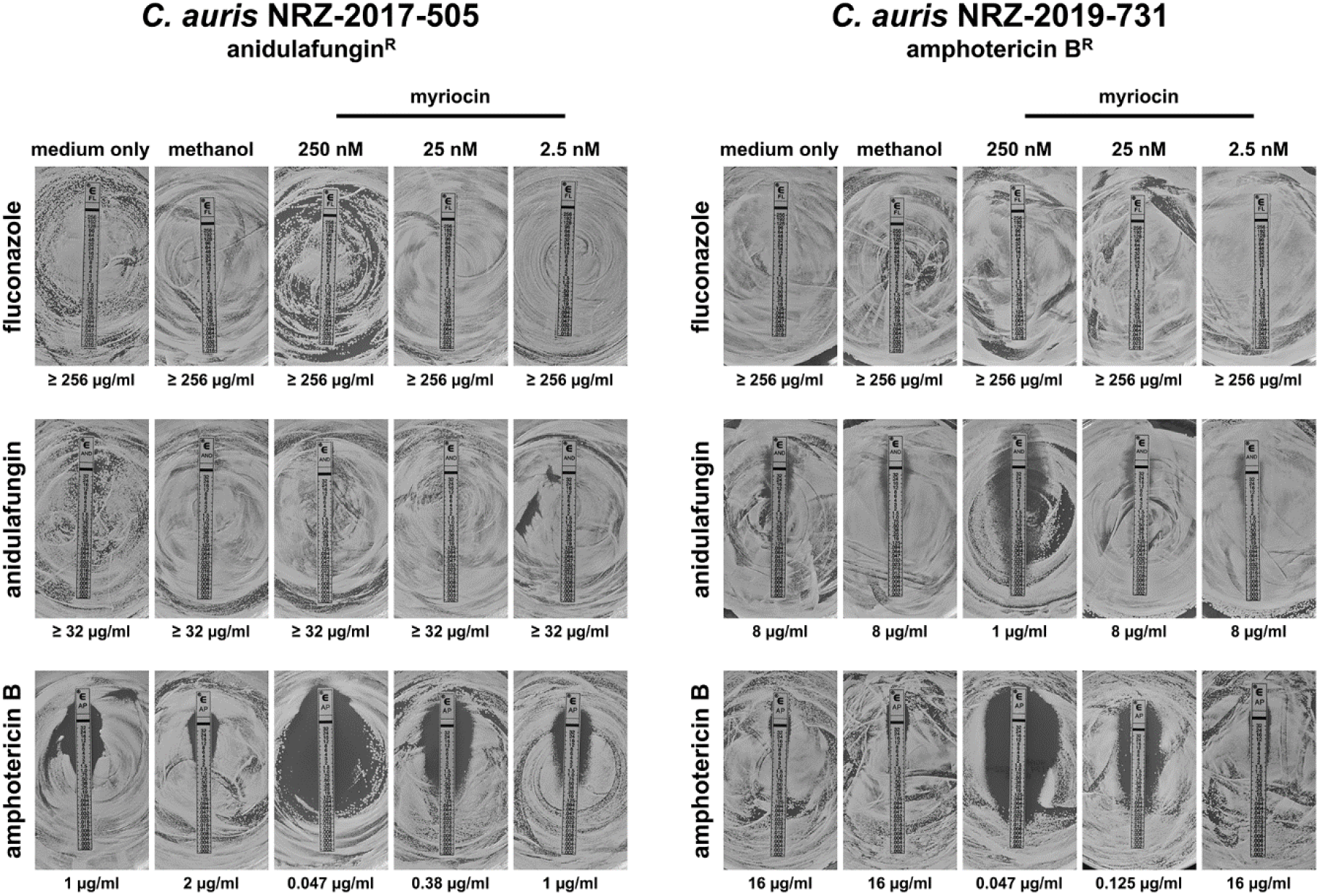
Addition of myriocin increases antifungal drug susceptibility of resistant *C. auris* strains. Cells of the *C. auris* strains NRZ-2017-505 and NRZ-2019-731 were plated onto RPMI1640 agar with or without the indicated concentrations of myriocin or methanol as solvent control. The indicated E-tests were applied, and plates were incubated at 35°C for 48 h before pictures were taken.

The isolate NRZ-2019-731 shows phenotypical resistance to amphotericin B with a MIC of 16 µg/ml on RPMI medium with or without methanol (Figure 3). Although no *FKS* hotspot mutation could be detected, the strain displayed an elevated MIC-value for anidulafungin of 8 µg/ml (Figure 3). Addition of myriocin had no effects on the susceptibility to fluconazole, but the highest concentration of the inhibitor (250 nM) caused an anidulafungin MIC-value decrease from 8 to 1 µg/ml (Figure 3). NRZ-2019-731 became susceptible to amphotericin B in presence of 250 and 25 nM myriocin as the MICs decreased from 16 µg/ml to 0.047 and 0.125 µg/ml (Figure 3). 2.5 nM seemed to be not sufficient to increase the susceptibility to the antifungal (Figure 3).

### Large scale screening for combinatory effects of myriocin and amphotericin B

Due to these findings, we further screened a total of 59 *C. auris* strains including primary and secondary isolates and CDC reference strains for the four major *C. auris* clades. We first tested their susceptibility against myriocin in a broth microdilution growth assay. With elevated myriocin concentrations an increasing number of impaired growing *C. auris* strains were observed. At 25 nM myriocin, all strains could grow. At 250 nM, four strains were no longer able to grow. Only 30 of the 59 tested strains could grow at 1 µM myriocin and this number further decreased to 15 at 2 µM myriocin (Figure 4 A). These data suggest that high susceptibility of *C. auris* to myriocin is a broad phenomenon and not a strain-specific one (Figure 4 A). In a second experiment all 55 strains were applied to a broth microdilution assay with amphotericin B alone or in combination with or 25 nM or 250 nM myriocin. Strains which did not grow at 250 nM myriocin were excluded (Figure 4 B). In presence of amphotericin B only, most of the strains had a MIC between 1 and 2 µg/ml. The latter was also the highest MIC which could be detected for any strain in this setting (Figure 4 B). Addition of 25 nM myriocin to amphotericin B lowered the MIC range most strains between 0.125 and 0.5 µg/ml (Figure 4 B). This effect was increased for the combination of amphotericin B and 250 nM myriocin as most of the strains had an MIC in the range between 0.03 and 0.125 µg/ml (Figure 4 B). None of the strains were able to grow at higher amphotericin concentrations than 0.5 µg/ml in presence of 250 nM myriocin (Figure 4 B).

**Figure 4.**
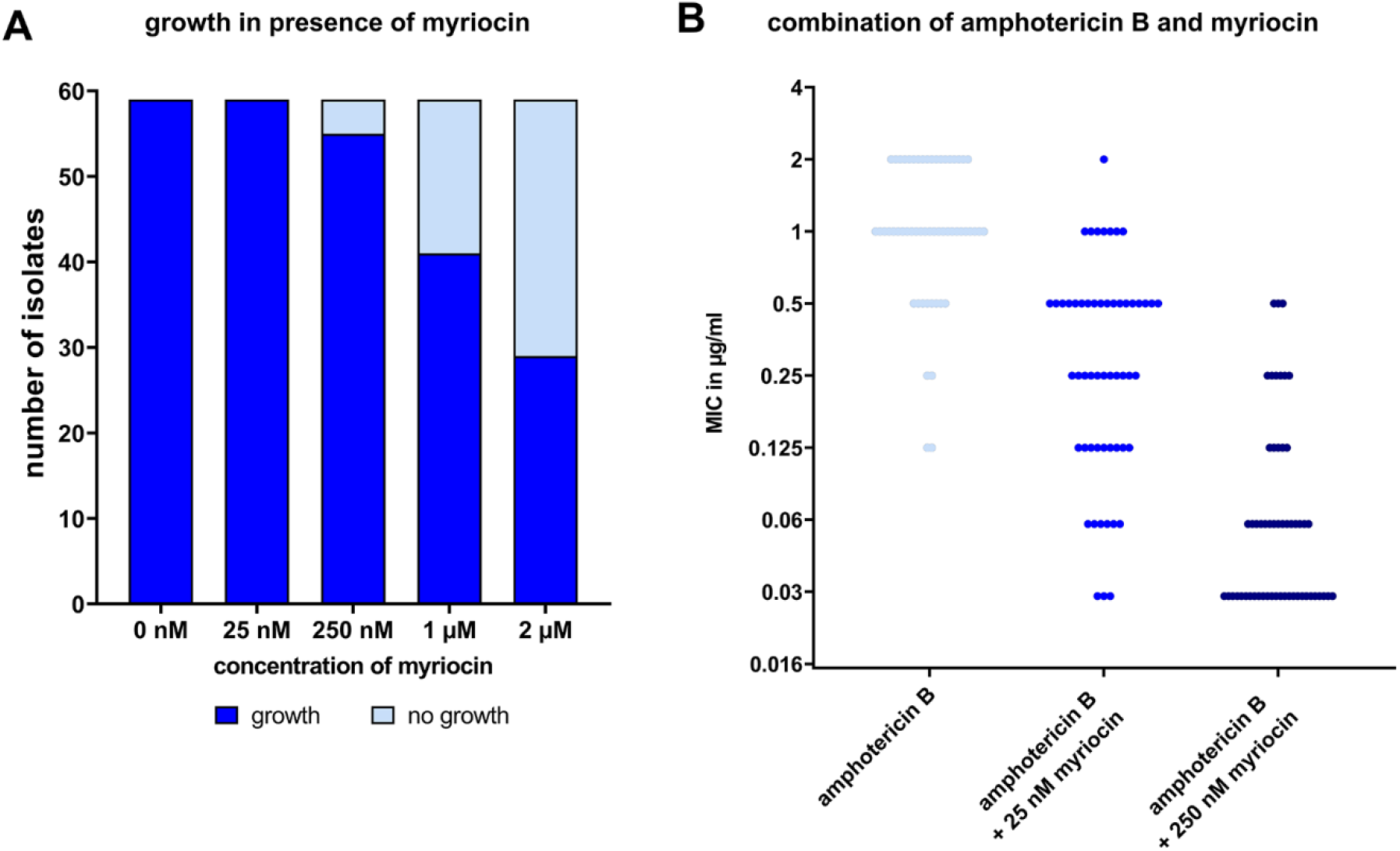
Susceptibility of *C. auris* to amphotericin B is increased in presence of myriocin in a large-scale screening. (A) A library of 59 *C. auris* strains were grown in a broth microdilution assay with RPMI1640 medium and the indicated myriocin concentrations for 24 h at 35°C. Wells without any sign of fungal growth were defined as “no growth”, all other wells were defined as “growth” even if there only single colonies visible. (B) A broth microdilution assay was performed with 55 *C. auris* library strains in presence of different concentrations of amphotericin B and myriocin. Plates were incubated for 24 h at 35°C before fungal growth was analyzed.

## Discussion

In accordance with previous works (15,24), we were able to show that all our tested *C. auris* isolates were highly susceptible to disruptions of the sphingolipid pathway by myriocin, a compound inhibiting the enzyme serine palmitoyl transferase in the initial step of *de novo* synthesis of sphingolipids (27). Therefore, we focused on the combination of myriocin with three different antifungals in this work. While the combination of myriocin and fluconazole had only limited synergistic effects, fungal susceptibility to anidulafungin and especially amphotericin B could be increased by the addition of the inhibitor. However, in the case of strains where phenotypic anidulafungin resistance is combined with *FKS* hotspot mutations, myriocin did not increase susceptibility to the antifungal drug. In contrast, even amphotericin B-resistant strains became susceptible to this polyene after the addition of sublethal myriocin concentrations. This indicates that sphingolipids play an important role in the ability of *C. auris* to handle amphotericin B. Together with the different sphingolipid composition (22-24), this could explain why some many clinical isolates of *C. auris* show a high tolerance to amphotericin B which is not common for *C. albicans or C. glabrata* (13,14,28). The antifungal activity of amphotericin B depends on the balance of different interactions with ergosterol, sphingolipids and phospholipids. A change of the sphingolipid composition in the cell membrane was shown to have direct influences on amphotericin B with other membrane components (29). It was already shown for *C. albicans* and *Saccharomyces cerevisiae* that a disruption of the sphingolipid biosynthesis by either inhibitors or gene deletion increases the susceptibility to amphotericin B (21 30, 31). A combination of sphingolipid synthesis inhibitors and fluconazole was shown to increase antifungal susceptibility of *C. albcians* and *C. glabrata* (32). It was also shown that inhibition of the *de novo* synthesis of sphingolipids in *C. albicans* abrogated biofilm formation and could lead to abnormal hyphal growth (31,33,34). Not much is known about the transcriptional control of the sphingolipid homeostasis in *Candida* species, but transcription factors Rtg1/3 might be involved, at least in *C. albicans* (35).

In contrast, inhibition of the *de novo* synthesis of sphingolipids by myriocin likely has no relevant effect on the *in vitro* susceptibility to azoles. Similar to our observations for *C. auris*, addition of the inhibitor had no negative effects on the susceptibility of *Aspergillus fumigatus* to azoles (36). However, sphingolipids might contribute to echinocandin-Fks interaction in *C. glabrata. C*hanges in their composition caused by addition of sphingosine might contribute to reduced echinocandin susceptibility in strains without *FKS* hotspot mutations (37). In contrast to our work with *C. auris*, addition of myriocin did cause increases in the susceptibility to anidulafungin in *C. glabrata* (37).

Together with previous findings (15, 21, 24, 30-32), our data suggest that a disruption of the sphingolipid pathway in *C. auris* could be an interesting approach for adjunctive antifungal therapies. This was already discussed for fungal sphingolipids at all (27). However, it should also be noted that a treatment with myriocin might have several side effects on the host (30). Therefore, myriocin itself might not be suitable for therapeutic usage but could be a base for the development of future antifungal drugs.

## Material and Methods

### Strains and media

All *Candida* sp. strains used in this study are listed in supplemental table 1. They were routinely maintained in YPD medium (20 g/l glucose, 20 g/l peptone, 10 g/l yeast extract, with 20 g/l agar if required) at 37°C.

### Growth tests with myriocin

2 mg myriocin powder (from Sigma Aldrich) were resuspend in 1ml methanol at 37°C to create a 5 mM stock solution which was routinely stored at -20°C. Selected *Candida* strains were grown overnight in YPD at 37°C. 5 µl of these overnight cultures were dropped in different dilutions onto YPD, RPMI1640 or SDG plates which contained 1 µM and 5 µM myriocin. Methanol was routinely used as solvent control. The plates were then incubated for 2 days at 30°C or 37°C.

### Antifungal Drug susceptibility testing

Antifungal drug susceptibility testing for single strains was performed with E-tests (Biomerieux) for amphotericin B, anidulafungin and fluconazole. *Candida* cultures were grown overnight in YPD at 37°C and 5×10^6^ cells/ml of these overnight cultures were plated onto RPMI1640 agar. E-test stripes were then applied according to the manufacturer’s instructions. To test the effects of myriocin, the chemical was plated on RPMI1640 agar in different concentrations ranging from 2.5 nM to 250 nM before the addition of fungal cells and application of the E-test stripe. The screening of the whole library of 59 clinical *C. auris* isolates was performed with broth microdilution similar to EUCAST protocols (38). In short, strains were grown in RPMI1640 medium with amphotericin B concentrations ranging from 0 to 8 µg/ml for 24 h at 37°C. If required, 25 nM or 250 nM myriocin were added to medium.

## Supporting information

The table contains all fungal strains which were used in this study.

## Acknowledgments

This project was in parts funded by the Deutsche Forschungsgemeinschaft (DFG GRK2581/1 to O.K.). We thank Anastasia Besenfelder, Ina Gaube and Sabrina Speiser for their excellent technical support.

